# Runx/Cbfβ regulates the development of tolerogenic Thetis cells

**DOI:** 10.1101/2025.08.27.672523

**Authors:** Chihiro Ogawa, Chengcheng Zou, Yoselin A Paucar Iza, Motoi Yamashita, Tyler Park, Junji Harada, Naoko Satoh-Takayama, Masahiko Kuroda, Chrysothemis C Brown, Ichiro Taniuchi

## Abstract

Establishing immune tolerance to gut microbiota and food antigens upon first exposures during early life is essential to prevent inflammatory bowel diseases and food allergy and depends on induction of peripherally induced Rorγt expressing regulatory T (Rorγt^+^ pTreg) cells^1, 2, 3, 4, 5, 6^. Recent studies have identified a critical role for Rorγt expressing antigen-presenting cells (APC), Thetis cells (TCs), in peripheral regulatory T (pTreg) cell differentiation and tolerance to food and commensal microbes^7, 8, 9, 10, 11^. TCs encompass four distinct subsets, and a subset of TCs, TC IV induces pTreg differentiation, but the transcription factors that control their differentiation are not fully known. Here, using orthogonal genetic approaches to impair Runx/Cbfβ activity, we show that development of specific TCs subsets is regulated by Runx/Cbfβ transcriptional factor complexes. While attenuated Runx3 by germline mutations resulted in a severe reduction of all Rorγt^+^ APCs, mice lacking one of two Cbfβ splicing variants, Cbfβ2, exhibited a loss of TC II, III and TC IV subsets with associated loss of Rorγt^+^ pTreg cells. Conditional inactivation of *Runx1* and *Runx3* genes by CD11c-Cre led to a specific loss of TC III and TC IV subsets. Strikingly, *CD11c-Cre* driven transgenic Runx expression, particularly Runx1, led to enhanced TC IV differentiation and thus Rorγt^+^ pTreg cells. Furthermore, transgenic Cbfβ2 by *CD11c-Cre* recovered only TC IV subset with restoration of Rorγt^+^ pTreg in Cbfβ2-deficient mice. Collectively, our findings establish a critical pathway for TC IV differentiation and provide new insights into therapeutic interventions to promote Rorγt^+^ pTreg induction in autoimmune and inflammatory diseases.

---

Establishing and maintaining tolerogenic immune responses to innocuous foreign antigens is vital for human health. Foxp3 expressing regulatory T cells (Tregs) are essential for immune tolerance and are first developed in the thymus, where self-antigens, such as Tissues Specific Antigens (TSA) presented by autoimmune regulator (Aire) expressing medulla thymic epithelium cells (mTEC) guides some autoreactive T cells to differentiate into Tregs^12, 13, 14, 15^. Generation of peripheral induced Foxp3^+^ Treg (pTreg) cells, in particular Rorγt expressing pTreg cells (Rorγt^+^ pTregs), from naive CD4^+^ T cells upon encounter with antigens derived from commensal microbiome and food is a key to establishing immune tolerance in the gut^3, 4, 5, 6^, and requires T cell priming by tolerogenic antigen-presenting cells (APCs)^16^. Several studies have established a new paradigm for intestinal tolerance, demonstrating the existence of a novel lineage of Rorγt^+^ antigen-presenting cells, Thetis cells (TCs), that encompass four distinct subsets. Among these, integrin αvβ8 expressing TCs, in particular TC IV, play a critical role in promoting microbiota^7^ and food-specific^17^ pTreg cells. Different genetic approaches have been used to narrow down the exact TC subset responsible for pTreg differentiation. These studies have shown that the tolerogenic APC resides within αvβ8 expressing TCs that lack Aire expression, suggesting that Aire^−^ αvβ8^+^ TC IV cells are the tolerogenic subset ^7, 10, 11, 17^. Other studies have highlighted a role for the transcription factors Prdm16 and Rorγt in the regulation of particular TC subsets^10, 11^ ; however the hierarchy of transcription factors that regulate TC development is not known. Intriguingly, *Runx3* gene inactivation in hematopoietic^18^ or CD11c-expressing cells^19^ resulted in spontaneous colitis development, in line with the genetic association of Runx3 in human inflammatory bowel disease (IBD)^20^. However, Runx/Cbfβ transcriptional heterodimer complexes have pleiotropic effects on immune cell differentiation including thymic-derived Tregs^21, 22^, effector CD4^+^ T cells^23, 24^, ILC3s/LTi cells^25, 26^ and classical dendritic cells (cDCs)^19^ and the cellular mechanisms underlying Runx/Cbfβ regulation of intestinal tolerance have not been established. Here, we set out to investigate the role of Runx/Cbfβ in Thetis cell differentiation.

## Colitis development and specific loss of Rorγt^+^ pTregs in Cbfβ2-deficient mice

Murine *Cbfb* gene produces two functional variants, Cbfβ1 and Cbfβ2, by alternative RNA splicing using different splicing donor signals within the exon 5. We previously generated a mouse line that specifically lacks the Cbfβ2 variant due to mutations, referred to as *Cbfb*^*2m/2m*27^. In addition to airway infiltration^27^, *Cbfb*^*2m/2m*^ mice spontaneously developed colitis at the age of three months (**Fig. 1a**) and histological changes appeared at around 8 weeks in *Cbfb*^*2m/2m*^ mice (**Extended Data Fig. 1a**). Given the critical role of Rorγt^+^ pTreg cells in gut microbiota tolerance^3, 4, 6^, we examined T cell subsets in small and large intestine lamina propria (SI-LP and LI-LP), and found that Rorγt^+^ pTregs were specifically lost in both SI-LP and LI-LP of *Cbfb*^*2m/2m*^ mice, while Rorγt^−^Foxp3^+^ Treg and Rorγt^+^Foxp3^−^ Th17 cells were not significantly changed (**Fig. 1b**). To determine the cell type responsible for impaired Rorγt^+^ pTreg cell differentiation in mice with germline *Cbfb*^*2m*^ mutation, we took advantage of *Rosa26*^*lsl-Cbfb2/+*^ mice^28^ to selectively rescue Cbfβ2 expression across a range of immune cell types. As expected, differentiation of Rorγt^+^ pTreg cells was restored in *Cbfb*^*2m/2m*^: *Rosa26*^*lsl-Cbfb2/+*^: *Vav1-iCre* mice (**Fig. 1c**). Restoration of Rorγt^+^ pTreg differentiation by transgenic Cbfβ1 expression (**Extended Data Fig. 1b-d**) suggested that low dosage of Cbfβ rather than loss of specific function of Cbfβ2 is causal for loss of Rorγt^+^ pTregs in *Cbfb*^*2m/2m*^ mice. By contrast, Rorγt^+^ pTreg cells were not rescued by transgenic Cbfβ2 expression in B or T lymphocytes in *Cbfb*^*2m/2m*^: *Rosa26*^*lsl-Cbfb2/+*^: *Mb1-* or *Cd4-Cre* mice respectively, indicating that impaired Rorγt^+^ pTreg cell differentiation in *Cbfb*^*2m/2m*^ mice occurs by T-cell-extrinsic mechanisms. Given the role of Rorγt^+^ APCs in pTreg differentiation, we next generated *Cbfb*^*2m/2m*^: *Rosa26*^*lsl-Cbfb2/+*^: *Rorc-Cre* and *Cbfb*^*2m/2m*^: *Rosa26*^*lsl-Cbfb2/+*^: *Cd11c-Cre* mice. While *Rorc-Cre* led to variable rescue in Rorγt^+^ pTreg cells, we observed complete restoration of Rorγt^+^ pTreg cell differentiation when transgenic Cbfβ2 expression was induced by *Cd11c-Cre* (**Fig. 1c** ), in line with previously reported loss of Rorγt^+^ pTregs by *Cbfb* inactivation by *Cd11c-Cre*^19^. Such variegated rescue by *Rorc-Cre* led us to wonder whether Cbfβ2 expression was required at an earlier stage of TC development before full acquisition of Rorγt expression. Given previous reports of IL7R-fate mapping in TCs^7^ and suggested descendancy from IL7R^+^ lymphoid progenitors^10^, we turned to *Cbfb*^*2m/2m*^: *Rosa26*^*lsl-Cbfb2/+*^: *Il7r-Cre* mice. In these mice, we observed complete restoration of Rorγt^+^ pTreg cells (**Fig. 1c**). Together, these observations indicate that a cell that has expressed CD11c, more specifically *Cd11c-Cre*, and IL7R during its developmental process requires Runx/Cbfβ2 for their development or pTreg-inducing function. Overall, these findings suggest a role for Runx/Cbfβ in Rorγt^+^ APC development.

**Figure 1.**
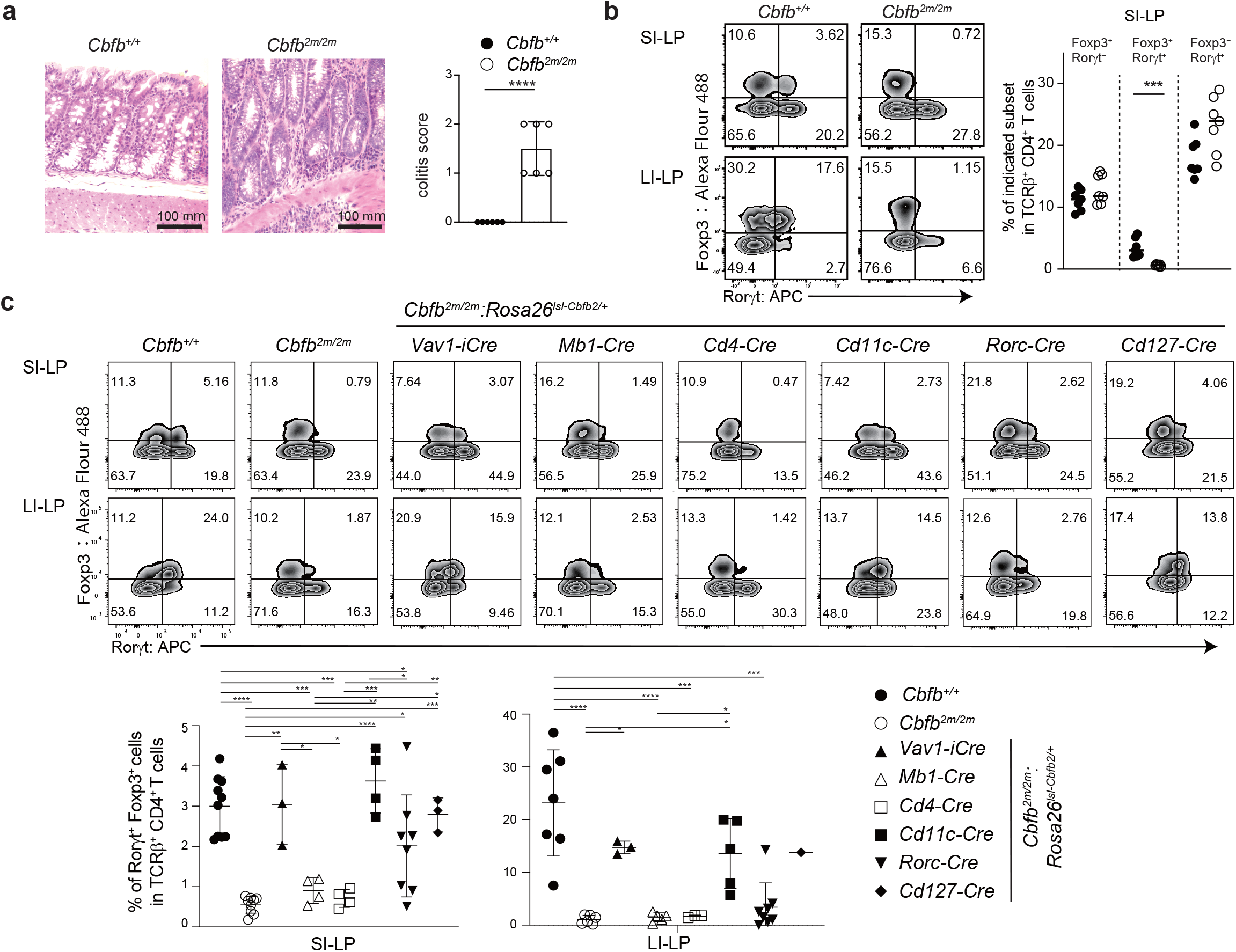
Cbfb^2m/2m^ice develop colitis due to dysfunction of antigen presenting cells. **(a)** Representative images of colon of *Cbfb*^*+/+*^ (*n*=6) and *Cbfb*^*2m/2m*^ mice (*n*=6). Graph at right showing a summary of colitis score. ****: *p* < 0.0001, unpaired *t*-test. Bars indicate 100 μm scale. **(b)** Zebra plots showing representative Rorγt and Foxp3 expression in CD4^+^ T cells of small intestine lamina propria (SI-LP) and large intestine lamina propria (LI-LP) of *Cbfb*^*+/+*^ (*n*=7) and *Cbfb*^*2m/2m*^ (*n*=7) mice. Graph showing statistical summary of the frequency of indicated cell subsets in CD4^+^ T cells. ***: *p* < 0.0005, unpaired *t*-test. **(c)** Zebra plots showing Rorγt and Foxp3 expression in CD4^+^ T cells of SILP and LILP of mice with indicated genotype; *Cbfb*^*+/+*^ (*n*≥ 7), *Cbfb*^*2m/2m*^ (*n*≥ 6), *Vav1-iCre* (*n*=3), *Mb1-Cre* (*n*≥ 4), *Cd4-Cre* (*n*≥ 3), *Cd11c-Cre* (*n*≥ 4), *Rorc-Cre* (*n*=8) and *Cd127-Cre* (*n*≥ 1). Transgenic Cbfb2 protein is expressed from the *Rosa26*^*lsl-Cbfb2*^ allele after excision of LoxP-Stop-LoxP (lsl) sequences by Cre-mediated recombination. Graph showing statistical summary of the frequency of Rorγt^+^Foxp3^+^ pTreg cells in SILP and LILP. For statistical analyses of LI-LP, *Cd127-Cre* model was not included since only one sample. *: *p* < 0.05, **: *p* < 0.001, ***: *p* < 0.0005, ****: *p* < 0.0001, One-way Anova analyses.

## Runx3/Cbfβ is required for TC development

To determine the role of Runx/Cbfβ in TCs differentiation, we examined Rorγt^+^ APC subset composition in mLNs of *Cbfb*^*+/+*^ and *Cbfb*^*2m/2m*^ mice. As previously shown^7^, TCs (CXCR6^−^ Rorγt^+^MHCII^+^ cells) can be divided into Ncam1^+^ TC I and Epcam^+^Ncam1^−^ cells which comprise CCR6^+^Nrp1^+^ TC II, CCR6^−^CD11c^+^CD11b^−^ TC III and CCR6^−^CD11c^+^CD11b^+^ TC IV subsets, and ILC3, which is also referred to as Lymphoid tissue inducer (LTi), are CXCR6^+^Rorγt^+^MHCII^+^ cells. Analysis of these Rorγt^+^ APC subsets revealed almost complete absence of TC II, III and IV subsets in *Cbfb*^*2m/2m*^ mice, but no change in TC I numbers (**Fig. 2a**). In addition, MHCII^+^ ILC3/LTi cell numbers were modestly impaired, in keeping with previous reports of Cbfβ2 regulation of LTi development^25^.

**Figure 2.**
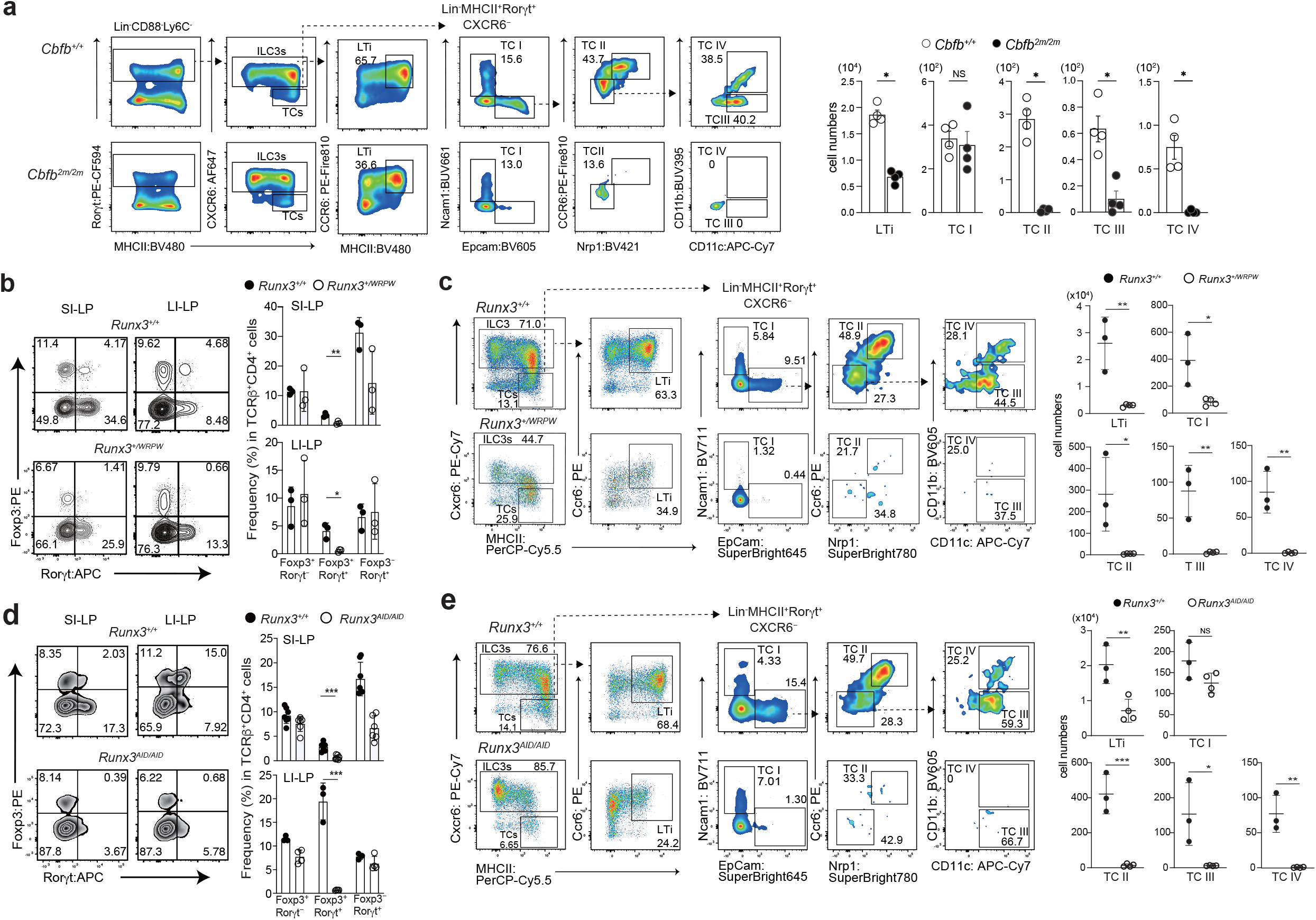
Severe reduction of Thetis cells in Cbfb^2m/2m^ and Runx3 mutant mice. **(a)** Representative flow cytometry showing gating of antigen presenting cell types in Rorγt^+^MHC-II^+^ population in mesenteric lymph nodes of two to three weeks-old *Cbfb*^*+/+*^ (*n*=4) and *Cbfb*^*2m/2m*^ (*n*=4) mice. Graphs showing statistical summary of numbers of indicated cell subsets. **(b)** Contour plots showing representative Rorγt and Foxp3 expression in CD4^+^ T cells of small intestine lamina propria (SI-LP) and large intestine lamina propria (LI-LP) of *Runx3*^*+/+*^ (*n*=3) and *Runx3*^*+/WRPW*^ (*n*=3) mice. (**c**) Representative flow cytometry showing gating of antigen presenting cell types in Rorγt^+^MHC-II^+^ population of mesenteric lymph nodes (mLNs) of *Runx3*^*+/+*^ (*n*=3) and *Runx3*^*+/WRPW*^ (*n*=4) mice. Graph showing a statistical summary of the frequency and numbers of Rorγt^+^MHC-II^+^ cells. **(d)** Zebra plots showing representative Rorγt and Foxp3 expression in CD4^+^ T cells of SI-LP and LI-LP of *Runx3*^*+/+*^ (*n*≥ 3) and *Runx3*^*AID/AID*^ (*n*≥ 4) mice. (**e**) Representative flow cytometry showing gating of antigen presenting cell types in Rorγt^+^MHC-II^+^ population of mLNs of *Runx3*^*+/+*^ (*n*=3) and *Runx3*^*AID/AID*^ (*n*=4) mice. Graph showing a statistical summary of the frequency and numbers of Rorγt^+^MHC-II^+^ cells. *: *p* < 0.01, **: *p* < 0.001, ***: *p* < 0.0005, unpaired *t*-test.

Cbfβ is a common partner for all three murine Runx family proteins, Runx1, Runx2 and Runx3. Among these, dysfunction of Runx3 has been shown to result in spontaneous colitis development^18^. To specifically address the role of Runx3 in intestinal tolerance, we generated a mouse line, *Runx3*^*WRPW*^, in which the terminal tyrosine (Y) residue within the WRPY motif in Runx3 was replaced with tryptophan (W). The WRPY motif is known to serve as a platform for recruiting transducin-like enhancer of split (TLE) co-repressor family protein^29, 30^. Since the WRPW motif in bHLH transcription factors, such as Hes1, has higher affinity to TLE family protein^31^, the Y to W replacement transforms Runx3 protein to a dominant repressor (manuscript submitted). Analysis of *Runx3*^*+/WRPW*^ mice revealed absence of Rorγt^+^ pTreg cells (**Fig. 2b**) with histological evidence of colitis by three months of age (**Extended Data Fig. 2a**). Although pLN and PP development is disrupted in *Runx3*^*+/WRPW*^ mice, we did not observe defects in the mLN of these mice. Importantly, numbers of Rorγt^+^MHC-II^+^ mLN cells in *Runx3*^*+/WRPW*^ mice were severely reduced (**Expanded Fig. 2b**). Accordingly, development of all TC subsets as well as LTi cells was almost completely abrogated in *Runx3*^*+/WRPW*^ mice, indicating that Runx3/Cbfβ complexes are responsible for the development of ILC3/LTi and TCs. However, since Runx3^WRPW^ mutant would interfere with Runx1 function when a Runx3^WRPW^-Runx1 heterodimer is formed (manuscript submitted), we cannot exclude possible redundant functions of Runx1 and Runx3 in TC development, analogous to the role of Runx proteins in LTi development^25^.

Thus, to further address the specific role of Runx3,we took advantage of another mouse model developed with the intention of inducible degradation of Runx3 by an Auxin-inducible degron 2 (AID2) system^32^, in which mAID degron tag sequences were inserted into endogenous Runx3 protein upstream of the C-terminal end (**Extended data Fig. 2c**). Unfortunately, such mAID degron tag insertion inadvertently led to impaired Runx3 function, as evidenced by absence of PP and most peripheral lymph nodes in *Runx3*^*AID/AID*^ mice. Using this mouse to study effects of hypomorphic Runx3, we found that pTreg generation was impaired (**Fig. 2d**), with development of colitis by three months of age in in *Runx3*^*AID/AID*^ mice (**Extended data Fig. 2d**). In addition, we observed absence of TC II, III and IV subsets with mild reduction of ILC3/LTi cells and no significant change in TC I numbers in *Runx3*^*AID/AID*^ mice (**Fig. 2e**). Collectively these data demonstrate a critical role for Runx3/Cbfβ complex in the development of TC II, III and IV subsets.

## Runx/Cbfβ regulates *Rorc* and *Prdm16* expression during TCs development

In our prior study, we identified a role for Runx/Cbfβ in the regulation of *Rorc* expression^27^. Given recent reports demonstrating dependence of Prdm16^+^Rorγt^+^ APCs on a +7kb *cis*-regulatory element in the *Rorc* locus^10, 11^, we wondered whether Runx/Cbfβ was required for Rorγt expression during TCs development. Among TCs subsets, we observed a spectrum of Rorγt expression with highest levels observed in TC II and low levels in TC I (**Extended Data Fig. 2e**). In addition, analysis of Prdm16 expression revealed its expression within TC II, III and IV (**Extended Data Fig. 2e**), but only low levels by a fraction of TC I, consistent with published scRNA-seq results^7, 10, 33^. Intriguingly, although TC I numbers were not impacted in *Cbfb*^*2m/2m*^ mice, the level of Rorγt expression was reduced (**Extended Data Fig. 2f**), suggesting a role for Runx/Cbfβ in regulation of *Rorc* expression in TCs. By contrast to the modest decrease in Rorγt expression in TC I, we observed complete absence of Prdm16 expressing TC I in *Cbfb*^*2m/2m*^ mice (**Extended Data Fig. 2f)**. The frequency of Prdm16^+^ TC I was also reduced in *Runx3*^*AID/AID*^ mice (**Extended Data Fig. 2f**). These observations suggested additional roles for Runx3/Cbfβ in regulation of *Prdm16* expression, presumably independent of Rorγt.

We previously identified a +11kb genomic region in the *Rorc* locus that was bound by Runx/Cbfβ complexes in fetal liver and required for Rorγt expression in fetal liver LTi cells^27^. Analysis of previously published single cell ATAC-seq data for TCs and ILC3s^7^ revealed accessible chromatin at the +11kb enhancer in all TCs subsets (**Fig. 3a**), as well as accessibility at the +7kb region in TC II-IV. To determine the role of *Rorc* +11kb enhancer (+11E) in regulating TC differentiation we analyzed +11E-deficient *Rorc*^Δ*+11E/*Δ*+11E*^ mice^27^. This revealed almost complete absence of both Rorγt^+^ pTreg and Rorγt^+^ Th17 cells in small and large intestine of *Rorc*^Δ*+11E/*Δ*+11E*^ mice (**Fig. 3b**). In line with the role of +11kb enhancer in LTi development, there were no lymph nodes or Peyer’s patches (PP) in these mice, making it impossible to examine TCs which predominantly reside in lymph nodes. To circumvent this, we generated fetal liver chimeric mice by transferring fetal liver from E14.5 *Rorc*^Δ*+11E/*Δ*+11E*^, knock in/knock out *Rorc*^*gfp/gfp*^, *Cbfb*^*+/+*^ or *Cbfb*^*2m/2m*^ embryos into sub-lethally irradiated recipient mice. Two weeks later, we observed reconstitution of TCs from wild-type *Cbfb*^*+/+*^ fetal liver but significantly impaired TCs numbers in *Cbfb*^*2m/2m*^ recipients and complete absence of TCs in *Rorc*^Δ*+11E/*Δ*+11E*^ and *Rorc*^*gfp/gfp*^ fetal liver recipients (**Fig. 3c**). Together, these observations identify a critical role for Runx/Cbfβ in TC II-IV development as well as in regulation of Rorγt expression at least in part through activation of +11E in the *Rorc* locus.

**Figure 3.**
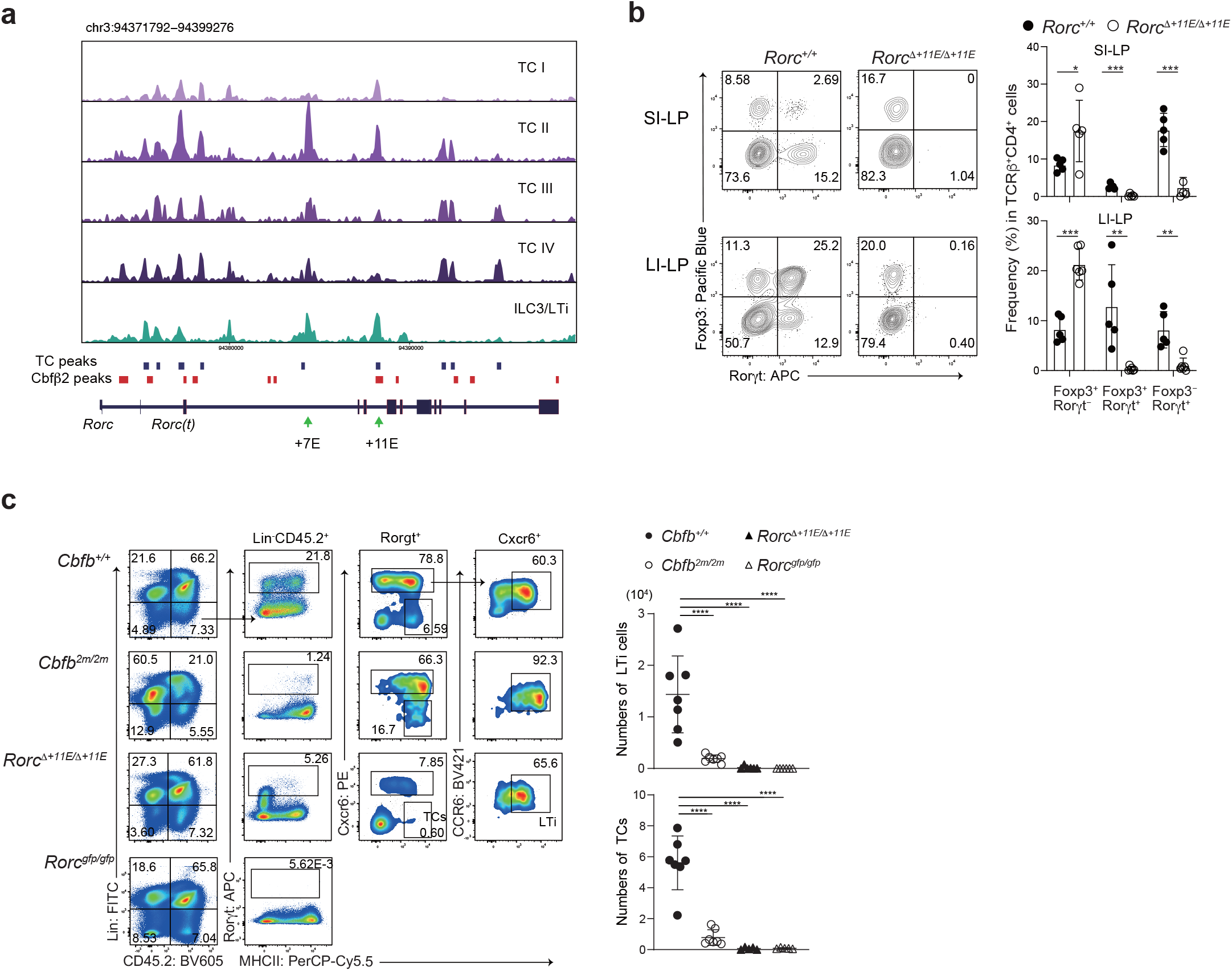
Lack of Thetis cell progenitors in Cbfb^2m/2m^ and Rorc^ΔE11/ΔE11^ice. **(a)** scATAC-seq track showing chromatin accessibility at the *Rorc* locus in indicated cell subsets. Cbfβ2 peaks detected by ChIP-seq are shown as red boxes. **(b)** Contour plots showing representative Rorγt and Foxp3 expression in CD4^+^ T cells of small intestine lamina propria (SI-LP) and large intestine lamina propria (LI-LP) of *Rorc*^*+/+*^ (*n*=5) and *Rorc*^Δ*E11/*Δ*E11*^ (*n*=5) mice. Graphs showing statistical summary of the frequency of indicated cell subsets in CD4^+^ T cells. *: *p* < 0.01, **: *p* < 0.005, ***: *p* < 0.0005, unpaired *t*-test. (**c**) Reconstitution of Rorγt^+^MHC-II^+^ cells from fetal liver cells (*n* ≥ 6 for each indicated genotype). 2 weeks after injection of fetal liver cells from CD45.2^+^ 14.5 dpc embryo with indicated genotype, mesenteric lymph nodes of CD45.1^+^ sub-lethally irradiated recipients were examined by flow cytometry. Representative pseudo-color plots showing Lin and CD45.2 expression, and Rorγt and MHC-II expression in Lin^−^CD45.2^+^ population, and Cxcr6 and MHC-II expression in Lin^−^CD45.2^+^ Rorγt^+^ cells, and CCR6 and MHC-II expression in Lin^−^CD45.2^+^ Rorγt^+^ Cxcr6^+^ cells. *: *p* < 0.05, ***: *p* < 0.0005, ****: *p* < 0.0001, One-way Anova analyses.

## Overexpression of Runx enhances TC differentiation and pTreg cell abundance

In contrast to the loss of Rorγt^+^ pTreg cells with germline hypomorphic *Runx3* mutation, *Runx3* inactivation by *Cd11c-Cre* did not lead to significantly reduction of Rorgt^+^ pTreg cells (**Extended Data Fig. 3a)**. Given redundant functions between Runx1 and Runx3 in other cell types such as cDC2s^19^, Runx1 may compensate Runx3 function for TCs development from a certain developmental stage where *Cd11c-Cre* was expressed. Indeed, differentiation of both Rorγt^+^ pTreg and Rorγt^+^ Th17 cells was severely inhibited in *Runx1*^*fl/fl*^: *Runx3*^*fl/fl*^: *Cd11c-Cre (Rx1/3*^*fl/fl*^: *Cd11c-Cre)* mice (**Fig. 4a)**. To clarify which Rorγt^+^MHC-II^+^ cell types have a history of *Cd11c-Cre* expression, we conducted lineage-tracing using a *Rosa26*^*lsl-tdTomato*^ reporter and found that almost all TC II, III and IV subsets were marked by tdTomato expression by *Cd11c-Cre*, while less than 40% of TC I and a minimal proportion of LTi cells were marked with tdTomato (**Fig. 4b**). In line with this observation, numbers of TC III and TC IV subsets were clearly decreased in *Rx1/3*^*fl/fl*^: *Cd11c-Cre* mice, although TC II numbers were not significantly reduced (**Fig. 4c**).

**Figure 4.**
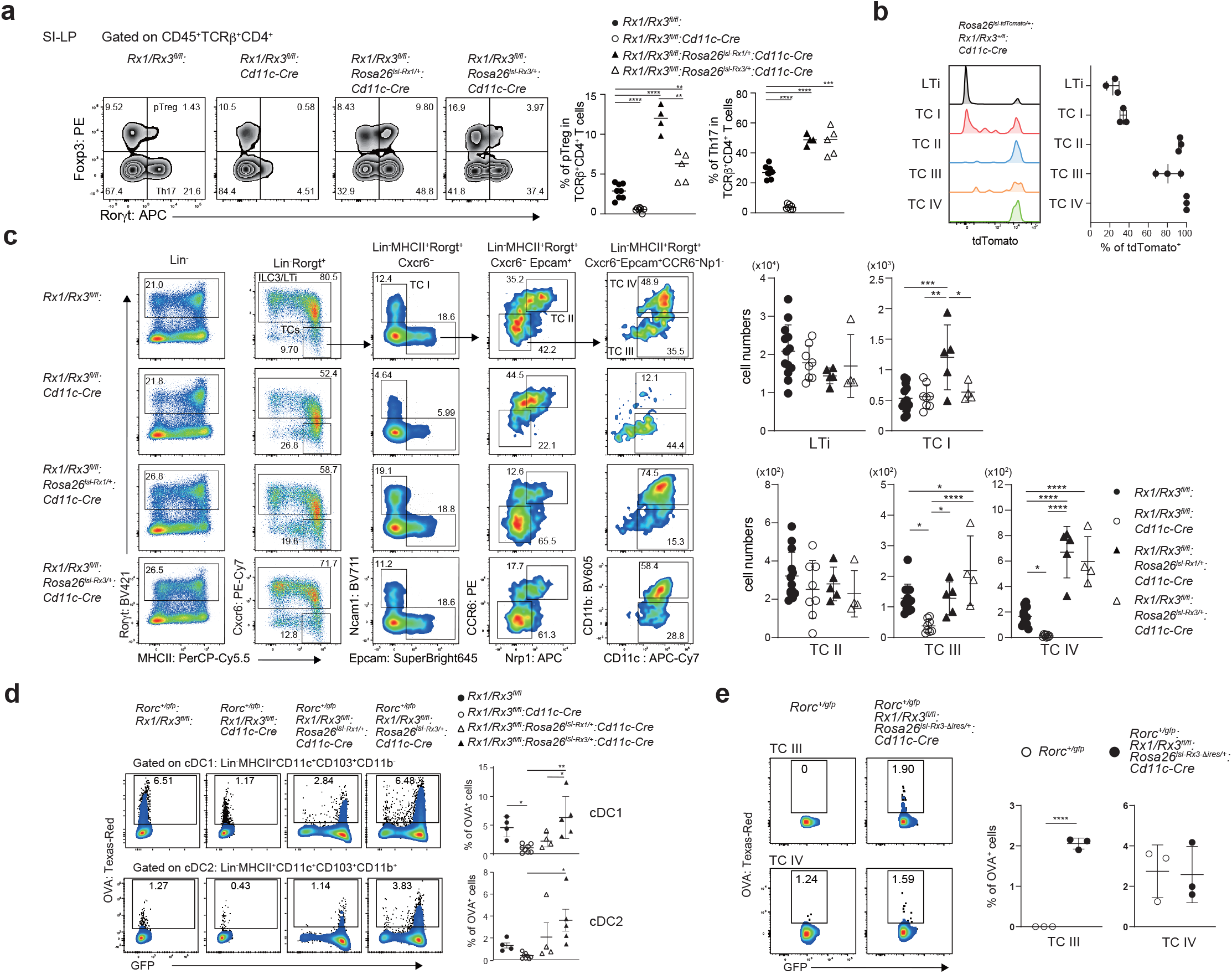
Runx/Cbfb regulates the development and function of TC III and TC IV. **(a)** Zebra plots showing representative Rorγt and Foxp3 expression in CD4^+^ T cells of small intestine lamina propria of mice with indicated genotype (*n* ≥4 for each indicated genotype). Graph at right showing a statistical summary. (**b**) Histograms showing tdTomato expression from the *Rosa26*^*lsl-tdTomato*^ locus in indicated cell subsets of *Rosa26*^*lsl-tdTomato/+*^*:Rx1/Rx3*^*+/fl*^*:Cd11c-Cre* mice (*n*=3). (**c**) Flow cytometry showing gating of antigen presenting cell types in Rorγt^+^MHC-II^+^ population in mesenteric lymph nodes of two to three weeks-old mice with indicated genotype (*n*≥4 for each indicated genotype). Graph at the right showing a statistical summary of numbers of LTi/ILC3 and each TC subset. (**d, e**) Representative pseudocolor plots showing TexasRed fluorescence in cDC1 and cDC2 (**c**) and TC III and TC IV (**d**) from mLNs 16 hours after oral gavage of OVA-TexasRed into two to three weeks of mice with indicated genotype (*n*≥4 for each indicated genotype) (**d**) and *Rorc*^*+/gfp*^ and *Rorc*^*+/gfp*^: *Rx1/3*^*fl/fl*^: *Rosa26*^*lsl-Rx3-*Δ*ires*/+^*:Cd11c-Cre* mice (n=3 for each indicated genotype) (**e**). Graphs at right showing a statistical summary. *: *p* < 0.05, **: *p* < 0.01, ***: *p* < 0.0005, ****: *p* < 0.0001, One-way Anova analyses (**d**) and unpaired *t*-test (**e**).

A major question is whether the Rorγt^+^ pTreg inducing ability of TCs can be harnessed for therapeutic approaches in intestinal inflammation such as IBD or food allergy. Given a rescue of Rorγt^+^ pTreg by transgenic Cbfβ2 expression, we wondered whether we could exploit Runx regulation to augment TC numbers and Rorγt^+^ pTreg differentiation. Taking advantage of having *Rosa26*^*lsl-Rx1*^ and *Rosa26*^*lsl-Rx3*^ mouse strains (manuscript submitted), we tested Runx1 and Runx3 ability by generating *Rx1/3*^*fl/fl*^: *Rosa26*^*lsl-Rx1/+*^: *Cd11c-Cre* and *Rx1/3*^*fl/fl*^: *Rosa26*^*lsl-Rx3/+*^: *Cd11c-Cre* mice. Both transgenic Runx1 and Runx3 restored, or rather enhanced, Rorγt^+^ pTreg development with a higher number of Rorγt^+^ pTreg cells by transgenic Runx1 (**Fig. 4a)**. This was associated with parallel changes in TCs, specifically within TC III and IV subsets. Severe abrogation of Th17 cell differentiation due to *Runx1/Runx3* inactivation by *CD11c-Cre* was also restored to levels above control mice with both transgenic Runx1 and Runx3 expression (**Fig. 4a)**. Underlying this phenomenon, we observed severe reduction of CD103^+^CD11b^+^ cDC2s, a critical cell type that has been implicated in intestinal Th17 cell differentiation due to their IL23 production capacity^34, 35^, was restored upon rescue of Runx1 or Runx3 expression (**Extended Data Fig. 3b**).

Having results showing rescue of specific TC subsets by *Cd11c-Cre* driven transgenic Runx protein, we also examined which TC subsets were restored by transgenic Cbfβ2 expression by *Cd11c-Cre* in *Cbfb*^*2m/2m*^ mice. Strikingly, only TC IV subset was restored in *Cbfb*^*2m/2m*^: *Rosa*^*lsl-b2/+*^: *Cd11c-Cre* mice (**Extended Data Fig. 4a**), indicating that recovery of TC IV alone was sufficient for restoration of Rorγt^+^ pTregs in *Cbfb*^*2m/2m*^ mice (**Fig. 1c**). Interestingly, we noticed that most of restored TC IV cells by transgenic Runx or Cbfβ2 lacked Prdm16 expression (**Extended Data Fig. 4b**). In contrast to the increased expression of Prdm16 among TC I with transgenic Runx1, transgenic Cbfβ2 did not impact Prdm16 expression by TC I (**Extended Data Fig. 4c**).

To further characterize function of Rorγt^+^MHC-II^+^ cell subsets, we next examined *in vivo* antigen uptake capacity using oral-gavage of TexasRed-Ova as well as a *Rorc*^*gfp*^ reporter allele. Sixteen hours after oral-gavage into *Rorc*^*+/gfp*^ mice, incorporation of TexasRed-Ova was detected in cDC1, cDC2 and TC IV with highest incorporation efficiency in cDC1 and TC IV, but not in ILC3/LTi neither TC I, II, III subsets (**Extended Data Fig. 4d**), as was recently reported^17^. Uptake of TexasRed-Ova by cDC1 and cDC2 was significantly reduced by loss of Runx1 and Runx3, and was restored by either transgenic Runx1 or Runx3 expression with more efficient restoration by Runx3 than Runx1 (**Fig. 4d**). After excision of *lsl s*equences, the *Rosa26*^*lsl-Rx3*^ locus produces GFP signal that overlayed Rorγt-GFP signals, preventing us to identify Rorγt^+^ cells. To overcome this experimental barrier, we generated *Rosa26*^*lsl-Rx3-*Δ*ires*^ allele by mutating *ires* sequences in the *Rosa26*^*lsl-Rx3*^ locus (**Extended Data Fig. 4e**) and confirmed that restored TC IV cells by transgenic Runx3 retained antigen-uptake capacity to similar extent to control TC IV cells (**Fig. 4e**). Interestingly, TC III subset acquired antigen-uptake capacity upon transgenic Runx3 expression (**Fig. 4e**). These observations suggested that Runx proteins regulate not only development but also function of cDCs and TCs.

## Discussion

In this work, we addressed the role of Runx/Cbfβ complexes in regulating Rorγt^+^ APC cells using different mouse lines. Our results not only provided definitive evidence that TC IV is the critical tolerogenic APC subset for Rorγt^+^ pTreg induction, but also shed new light on possible manipulation of TC IV development by engineering Runx expression. Our data suggest that attenuation of Runx3/Cbfβ complexes by germline mutation results in reduced activation of *cis*-regulatory elements, such as +7 and +11 kb enhancers, in the *Rorc* locus, leading to impaired initiation of TC developmental programs. As such, *Rorc-Cre* was not always activated in *Cbfb*^*2m/2m*^ mice during TCs development, causing a variegated rescue of Rorγt^+^ pTregs in *Cbfb*^*2m/2m*^: *Rosa26*^*lsl-Cbfb2/+*^: *Rorc-Cre* mice. Inactivation of *Prdm16* by *Rorc-cre* was shown to result in lack of tolDCs^11^ including TC II, III and IV. Loss of Prdm16^+^ TC I in *Cbfb*^*2m/2m*^ mice suggests that Runx/Cbfβ could be involved in *Prdm16* regulation. Emergence of TC IV lacking Prdm16 expression by transgenic Cbfβ2 or Runx protein suggest that Prdm16 is required for TC IV development only at early stage and becomes dispensable to maintain TC IV subset at later stage. There should be a time-lag of transgenic Runx protein induction after loss of endogenous Runx protein by *Cd11c-Cre*. Therefore, failure of Prdm16^+^ TC IV recovery by transgenic Runx protein suggest a narrow time window for recovery of *Prdm16* activation by Runx re-expression.

Distinct dependency on Runx/Cbfβ for differentiation of each TC subset highlighted the heterogeneity of TCs and suggest that each subset, which emerges during TC differentiation from as yet unidentified precursors, has a differential requirement for Runx/Cbfβ. Future studies, addressing the development pathways for individual TC subsets are required to further understand the role of Runx/Cbfβ in TC subset specification. Considering the essential role of TC IV in establishing tolerance to microbiome-derived and food antigen^11, 17^, understanding the transcriptional regulation of TC IV is crucial for insights into therapeutic possibilities in inflammatory bowel disease and food allergy. Our results showed that the development program of TC III and TC IV can be manipulated by exogenous Runx expression. In addition, Runx proteins play a role in regulating antigen uptake function of cDCs and TCs. It remains elusive how transgenic Runx1 induced by *Cd11c-Cre* enhances Rorγt^+^ pTreg differentiation. Besides the increase in number of TC IV, enhanced function of TC IV and other uncharacterized APCs could be involved in this mechanism. Unraveling mechanisms by which Runx exerts immunomodulatory activity during APCs differentiation is essential and may be applied to develop novel therapeutic approach for immune-related diseases through increase of Rorγt^+^ pTreg cells.

## Acknowledgments

We thank Ms Sawako Muroi for ES cell work, Ms. Noriko Yoza for the cell sorting, Mr. Yusuke Iizuka for genome editing, and Ms Hitomi Tatsumi and Ms Yurie Kawamoto for *in vitro* fertilization (IVF), Ms Yuria Taniguchi, Ms Chizuko Miyamoto, Sawako Muroi for genotyping and Ms Tomoko Kageyama for preparation of lamina propria cells. This work was supported by Grant-in-Aid for Scientific Research (C) (24K10277 to C.O.), Grant-in-Aid for Scientific Research (B) (17H04090 to I.T.), Grant-in-Aid for Scientific Research on Innovative Area (19H05747 to I.T.). C.C.B. was supported by National Institutes for Health National Institute of Allergy and Infectious Diseases (DP2 AI171116-01), the G. Harold and Leila Y. Mathers Foundation, a Pew Scholar award, W.M. Keck Foundation and Pershing Square Sohn Alliance. Y.P.I. was supported by an HHMI Gilliam Fellowship.

## Author contributions

C.O., C.Z., Y.P.-L., J.H. and N. S.-T. analyzed phenotypes of gene modified mice. M.Y performed TC reconstitution from fetal liver. T.P. analyzed scATAC-seq data. M.K. performed histological analyses. C.C.B. provided technical advice and wrote the manuscript. I.T. designed this the study, generated mice and wrote the manuscript.

## Additional Information

Supplementary Information is available for this paper. Correspondence and requests for materials should be addressed to Ichiro Taniuchi.

## Supplementary Materials

Materials and Methods

Extended Data Figures 1-4

Extended Table 1

## Online Methods

### Mice

*Cbfb*^*2m 27*^, *Rx1^flox 36^*, *Rx3^flox 36^*, *Mb1-Cre ^37^, CD4-Cre ^38^, CD11c-Cre ^39^*, *IL7Rα-Cre ^40^*, *Rosa26^lsl-Cbfb2 27^* and *Rorc^Δ+11E 27^* mice were previously described. *Vav1-iCre* (#018968), *Rorc-Cre* (#022791) and *Rosa26*^*tdTomato*^ (#007909) mice were purchased from Jackson laboratory. *Runx3*^*WRPW*^ mice will be described elsewhere (manuscript submitted). In brief, zygotes generated from C57BL/6NJcl strain purchased from CLEA Japan were injected with mRNA encoding humanized S. pyogenes Cas9, single guide RNA (sgRNA) and single strand donor DNA, both of which were synthesized at INTEGRATED DNA TECHNOLOGY (IDT), at animal facility at RIKEN IMS. *Runx3*^*AID*^ mice and *Rosa26*^*lsl-Rx3-*Δ*ires*^mice were also generated by genome editing conducted at RIKEN IMS. Sequences for sgRNA, donor DNA used in genome editing were listed in supplementary table 1. To construct the target vector for *R26*^*lsl-Cbfb1*^ mice, cDNA encoding Cbfβ1 protein was amplified by PCR to add AscI sites at both ends. This cDNA fragment was ligated into an AscI-cleaved pCTV vector (#15912, Addgene). 30 mg of the target vector was linearized by AsiSI enzyme and was transfected into the M1 ES cell line by electroporation as previously described ^27^. After G418 selection, G418 resistant ES clones were screened for homologous recombination event by PCR. Appropriate ES clones harboring *Rosa26*^*lsl-Cbfb1*^ allele were aggregated with blastocysts to generate chimera mice, through which the *Rosa26*^*lsl-Cbfb1*^ allele were germline transmitted to the offspring. *Rosa26*^*lsl-Rx1*^ and *Rosa26*^*lsl-Rx3*^ mice will be described elsewhere (manuscript submitted), but in brief, these mouse strains were also generated in a similar way to the generation of *R26*^*lsl-Cbf1*^ mice.

### Histology

Mouse intestinal tissues were harvested, fixed in 10% neutral buffered formalin, and paraffin embedded according to standard procedures. Four-micrometer sections were cut using a paraffin microtome with steel blades. Sections were stained with hematoxylin and eosin. Evaluation of inflammation in murine colitis was performed according to the previously described colitis score^41^.

### Tissue processing

Lamina propria lymphocytes (LPLs) from the small and large intestine were isolated as previously described^19^. The small intestines were cut into 5 mm pieces and incubated in 20 mL of RPMI containing 2% FBS, 5 mM EDTA, with shaking at 200 rpm and 37 °C for 20 min. After incubation, the tissues were washed with PBS twice by vortexing for 20 seconds to remove epithelial cells. The rest of the tissues were filtered and digested with RPMI including 2% FBS, 0.5mg/mL of collagenase IV (sigma C-5138), 50 μg/mL of DNase (043-26773; FUJIFILM) at 200 rpm and 37°C for 30 min. Digested tissues were passed through a 100 µm strainer and subjected to Percoll gradient centrifugation at 1800 rpm for 20 minutes using 40% and 80% Percoll in RPMI with 2% FBS. Cells located between the 40% and 80% Percoll layers were collected as LPLs.

For staining of Rorgt^+^MHC-II^+^ cells from mLNs, all mLNs were collected in RPMI containing 10% FBS, 0.5 mg/mL collagenase IV, and 50 μg/mL DNase I, and incubated at 37 °C for 30 minutes with shaking. After incubation, the digested samples were passed through a 100 μm cell strainer and centrifuged at 1200 rpm for 5 minutes at 4 °C.

### Flow cytometry

Surface molecules were stained with specific antibodies by incubating the cells for 30 minutes at room temperature for mLN cells, or for 10 minutes at room temperature for other cell types. The following surface marker antibodies were purchased from BD Biosciences, BioLegend, or Thermo Fisher Scientific: B220 (RA3-6B2), Ccr6 (29-2L17), CD4 (RM4-5), CD11b (M1/70), CD11c (N418), CD19 (6D5), CD24 (M1/69), CD45 (30-F11), CD45.1 (A20), CD45.2 (104), CD56 (809220), CD64 (X54-5/7.1), CD88 (20/70), CD103 (M290), CD304 (3E12), CD326 (G8.8), Cxcr6 (SA051D1), Gr-1 (RB6-8C5), NK1.1 (PK136), MHC-II (M5/114.15.2), Siglec-F (S17007L), TCRβ (H57-597), and γδTCR (GL3). For intracellular staining, cells were fixed and permeabilized using the Transcription Factor Buffer Set (562574, BD Biosciences) after surface staining. The following intracellular antibodies were purchased from BD Biosciences or Thermo Fisher Scientific: Foxp3 (FJK-16s) and Rorγt (B2D and Q31-378). For intracellular Prdm16 staining, a rabbit monoclonal anti-Prdm16 antibody (ab303534, Abcam) was used as the primary antibody, followed by a Goat anti-Rabbit IgG (H+L) Cross-Adsorbed Secondary Antibody, Alexa Fluor™ 647 (A21244, Invitrogen). Dead cells were excluded using the Fixable Viability Dye eFluor™ 506 (65-0866, Invitrogen). Multi-color flow cytometric analysis was performed using BD FACSCanto™ II and BD FACSAria™ Fusion (both from BD Biosciences), and data were analyzed with FlowJo™ software (BD Biosciences). Cell sorting was carried out using BD FACSAria™ III and BD FACSAria™ Fusion (BD Biosciences).

### Immuno-blotting

CD8^+^ T cells were isolated from the spleen using the EasySep™ Mouse CD8^+^ T Cell Isolation Kit (STEMCELL Technologies). Cell lysates from total thymocytes or CD8^+^ T cells were resolved by SDS-PAGE and transferred to Immobilon-P Transfer Membranes (Millipore). Membranes were probed with an appropriate primary antibody, rabbit anti-panCbfβ, anti-Cbfβ1, anti-Cbfβ2^27^, rabbit anti-Runx3 (D6E2, #9647, Cell Signaling), anti-Gapdh (6C5, sc-32233, Santa Cruz) and HRP-conjugated secondary antibody, and immune complexes were detected using ECL Prime (GE Healthcare) with Amersham Imager 600 (GE Healthcare).

### Fetal liver transfer

Fetal liver collected from 14.5 dpc CD45.2^+^ embryos was used to make single cell suspension. Recipient CD45.1^+^ C57BL/6N mice were irradiated at 9.5 Gy. The irradiated CD45.1^+^ C57BL/6N mice were injected with a half to whole fetal liver cells of CD45.2^+^ one embryo. Two to three weeks after injection, the recipient mice were analyzed by flowcytometry for reconstitution of Rorgt^+^MHC-II^+^ cells.

### Oral gavage and antigen uptake

To assess OVA uptake, P14-21 *Rorc*^*+/gfp*^ and *Rorc*^*+/gfp*^*:Rx1/3*^*fl/fl*^*:Cd11c-Cre* mice were gavaged with 200 μl of 20 mg/ml Texas-Red conjugated OVA (O23021, Invitrogen). 16 hours later, cells in mLNs were analyzed by flow cytometry.

### Cbfβ2 ChIP-seq analysis

FASTQ files of Cbfβ2 ChIP-seq data were obtained from GSE90794^27^. Reads were trimmed using TrimGalore v0.6.19, aligned to the mouse reference genome (GENCODE GRCm38 release M22) using bowtie2 v2.5.1, and deduplicated. Peaks were identified using MACS2 v2.2.7.1 (-f BAM -g mm --nomodel --pvalue 0.1 --shift 0 --extsize 165 --call-summits).

### scATAC-seq analysis

The scATAC-seq data were obtained from GSE205065^7^ and processed using ArchR v1.0.1 as described previously. Briefly, Latent Semantic Indexing (LSI) was performed using IterativeLSI function (iterations=10, varFeatures=100,000) of ArchR, and cells were clustered (method=Seurat, k.param = 30, resolution = 1.2) using 30 LSI components. For peak calling, clusters for similar cells were grouped: C1 (TC IV), C2-4 (TC I,II,III), C5-6 (NCR+ ILC3), and C7-13 (LTi). Peaks were called on each group using MACS2 v2.2.7.1 (--gsize mm --qval 0.01 --nomodel --ext 200 --shift -100 --call-summits). Peak summits were extended by 100 bp in each direction. Regions extending outside of mm10 chromosomes, arising from chrY or chrM, overlapping with blacklist regions precompiled by ArchR (merged from the ENCODE mm10 v2 blacklist regions from https://github.com/Boyle-Lab/Blacklist/blob/master/lists/mm10-blacklist.v2.bed.gz and mitochondrial regions that are highly mappable to the mm10 nuclear genome from https://github.com/caleblareau/mitoblacklist/blob/master/peaks/mm10_peaks.narrowPeak), or containing ‘N’ nucleotides (>0.001 of the sequence) were filtered. Regions from groups of TC clusters (C1 and C2-4) were compiled, and overlapping regions were merged to their union, resulting in a non-overlapping set of 101,264 peaks for the TC clusters. Genome tracks were visualized using plotBrowserTrack function in ArchR with peaks from TC clusters and Cbfb2 ChIP-seq.

### Statistical analyses

Statistical analysis was performed by Anova one-way analyses and unpaired *t*-test using GraphPad Prism (v8.4.3) (Graphpad software, CA).

## Data availability

The scATAC-seq data and Cbfβ2 ChIP-seq data supporting the findings of this study were obtained from the NCBI Gene Expression Omnibus (GEO) under accession numbers GSE205065 and GSE90794, respectively. For the scATAC-seq analysis, the reference mouse genome mm10 was used for mapping, while the GENCODE GRCm38 release M22 mouse reference genome was used for alignment in the Cbfβ2 ChIP-seq analysis. All other data are available from the corresponding authors upon reasonable request. Source data are provided with this paper.

## Legends for Extended Data Figures

**Extended Data Figure 1. Restoration of Ror**γ**t**^**+**^ **pTreg by transgenic Cbf**β**1 in Cbfb^2m/2m^ice**. (**a**) Representative images of large intestine of four, six and eight weeks old *Cbfb*^*2m/2m*^ mice. The right graph showing summary of colitis scores. (**b**) A scheme showing the structure of the *Rosa26* allele with cDNA fragment inserted for the inducible expression of transgenic Cbfβ1 after removal of loxP-Neo/STOP-loxP (lsl) sequences by Cre-mediated site-specific recombination. (**c**) One representative image of two immunoblot experiments showing Cbfβ1 expression in thymocytes of mice with indicated genotype. The graph at right showing numbers of Peyer’s patches (PPs) (*n*≥3 for each indicated genotype). ****: *p* < 0.0001, One-way Anova analyses. (**d**) Zebra plots showing representative Rorγt and Foxp3 expression in CD4^+^ T cells of small intestine lamina propria (SI-LP) of mice with indicated genotype (*n*≥4 for each indicated genotype). Graph showing a statistical summary of the frequency of indicated cell subsets in CD4^+^ T cells. *: p < 0.05, **: p < 0.01, unpaired *t*-test.

**Extended Data Figure 2. Colitis development by attenuated Runx3 function**.(**a**) Representative images of colon of *Runx3*^*+/+*^ (*n*=4), *Runx3*^*+/WRPW*^ (*n*=4) mice. Graph at right showing a summary of colitis score. Bars indicate 100 μm scale. (**b**) Representative pseudocolor plots showing Rorγt and MHC-II expression in Lin^−^CD64^−^Ly6c^−^ population of mesenteric lymph nodes (mLNs) of *Runx3*^*+/+*^ (*n*=6), *Runx3*^*+/WRPW*^ (*n*=4) and *Runx3*^*AID/AID*^ (*n*=4) mice. Graph showing absolute numbers of Rorγt^+^MHC-II^+^ cells. ***: p < 0.005, One-way Anova analyses. (**c**) A schematic structure of around exon 6 in murine *Runx3* gene and structure of donor ssDNA for insertion of mAID sequence. Amino acid sequences at the C-terminal of Runx3^AID^ mutant protein are shown; GS linker (green), mAID (red) and Runx3 derived amino acid (black). One representative result of two immunoblots with α-Runx3 antibody showing Runx3 and Runx3^AID^ protein expression in CD8^+^ T cells. Gapdh immunoblot was used as a loading control. (**d**) Representative images of colon of *Runx3*^*+/+*^ (*n*=5), *Runx3*^*AID/AID*^ (*n*=7) mice. Graph at right showing a summary of colitis score. Bars indicate 100 μm scale. (**e**) Histograms showing Rorγt and Prdm16 expression in indicated cell subsets. Mean fluorescent intensity (MFI) of Rorγt expression was indicated. (**f**) Left histograms showing Rorγt expression in TC I of *Cbfb*^*+/+*^ (*n*=4) and *Cbfb*^*2m/2m*^ (*n*=4) mice. Middle and right histograms showing Prdm16 expression in TC I subset of *Cbfb*^*+/+*^ (*n*=4), *Cbfb*^*2m/2m*^ (*n*=4) mice and *Runx3*^*+/+*^ (*n*=3) and *Runx3*^*AID/AID*^ (*n*=4) mice, respectively. Graph showing a statistical summary of Rorγt MFI (left) and the frequency of Prdm16^+^ TC-I. ** p < 0.01, ***: p < 0.005, ****: p < 0.0001, unpaired *t*-test.

**Extended Data Figure 3. Rescue of cDC2 development by transgenic Runx expression.**(**a**) Zebra plots showing representative Rorγt and Foxp3 expression in CD4^+^ T cells of small intestine lamina propria (SI-LP) of Rx3^*fl/fl*^ (*n*=3) and Rx3^*fl/fl*^: *Cd11c-Cre* (*n*=3) mice. Graph showing a summary of frequencies of Rorγt^+^ pTreg cells. (**b**) Contour plots showing CD103 and CD11b expression in Lin^−^MHC-II^+^CD11c^+^ cells in mLNs of mice with indicated genotype (*n*=4 for each indicated genotype). Graph showing a summary of frequencies of cDC1 (CD103^+^CD11b^−^) and cDC2 (CD103^+^CD11b^+^) cells. *: p < 0.05, ****: p < 0.0001, One-way Anova analyses.

**Extended Data Figure 4. Specific rescue of TC IV development by transgenic Cbf**β**2 induced by Cd11c-Cre**. (**a**) Representative flow cytometry showing gating of antigen presenting cell types in Rorγt^+^MHC-II^+^ population in mesenteric lymph nodes of two to three weeks-old *Cbfb*^*+/+*^ (*n*=5), *Cbfb*^*2m/2m*^ (*n*=5) and *Cbfb*^*2m/2m*^: *Rosa26*^*lsl-cbfb2/+*^: *Cd11c-Cre* (*n*=5) mice. Graphs showing statistical summary of numbers of indicated cell subsets. *: p < 0.05, ***: p < 0.005, ****: p < 0.0001, One-way Anova analyses. (**b, c**) Histograms showing Prdm16 expression in TC IV (**b**) and TC I (**c**) subsets of 2-3 weeks old mice with indicated genotype (*n*≥3 for each indicated genotype). Figures in histograms showing the frequency of Prdm16 expressing cells of control mice. Graph showing summary of the frequency of Prdm16^+^ TC IV (**b**) and TC I (**c**). *: p < 0.05, **: p < 0.01, ***: p < 0.005, One-way Anova analyses. (**d**) Representative pseudocolor plots showing Texas-Red fluorescence in indicated cell subset from mLNs 16 hours after oral gavage of OVA-Texas-Red into three-week-old *Rorc*^*+/gfp*^ mice (*n*=3). (**e**) A scheme showing the structure of the *Rosa26*^*lsl-Rx3*^ allele and strategy for targeting mutations into the *ires* sequences by CRISPR/Cas9 genome editing with two sgRNAs. Sequence of non-functional *ires* after genome editing is shown in which sequences deleted were highlighted with yellow color font. gRNA sequences are indicated with wavy underline.

