## Supplementary figures and images for "Runx/Cbfβ regulates the development of tolerogenic Thetis cells"

### Extended Figures

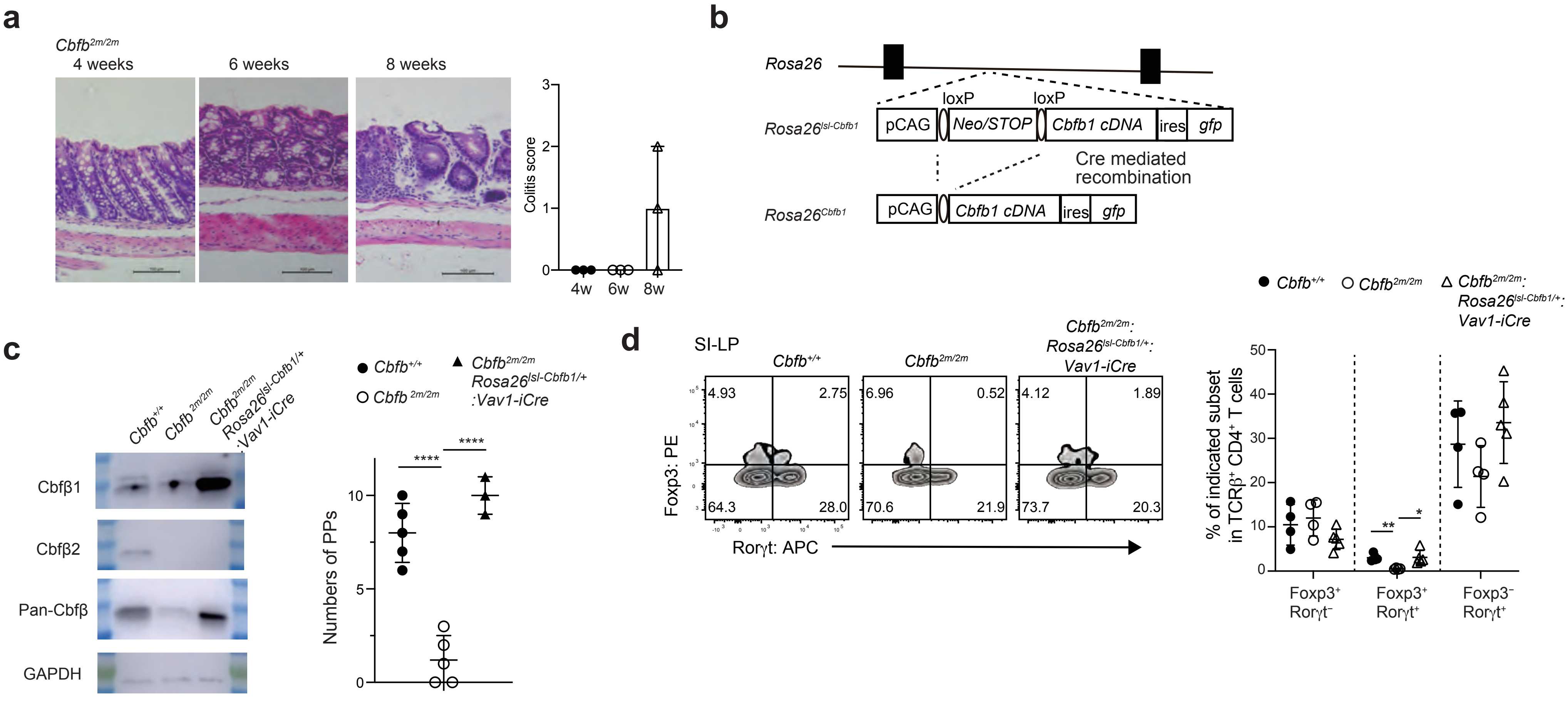


**Extended Data Figure 1**

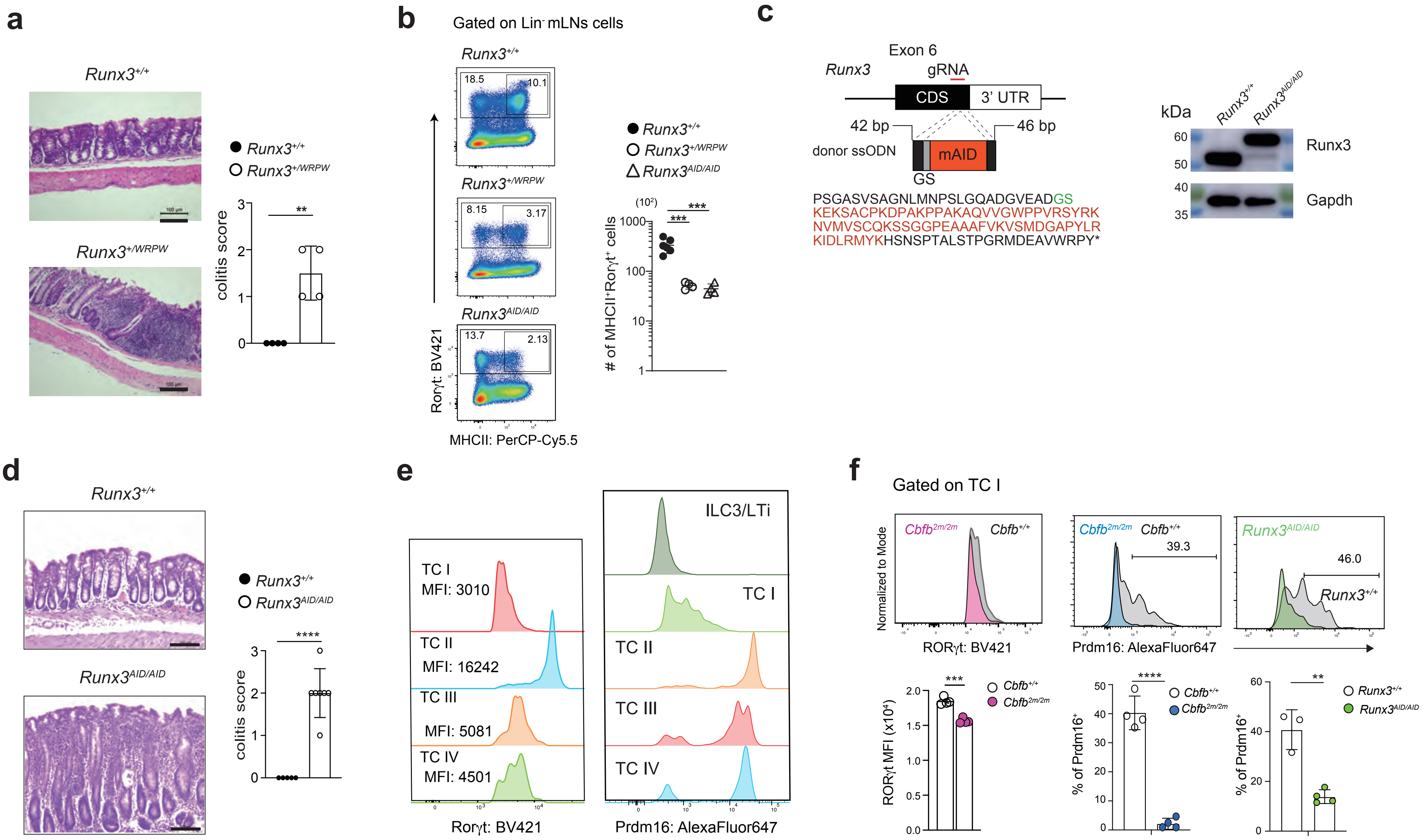


**Extended Data Figure 2**

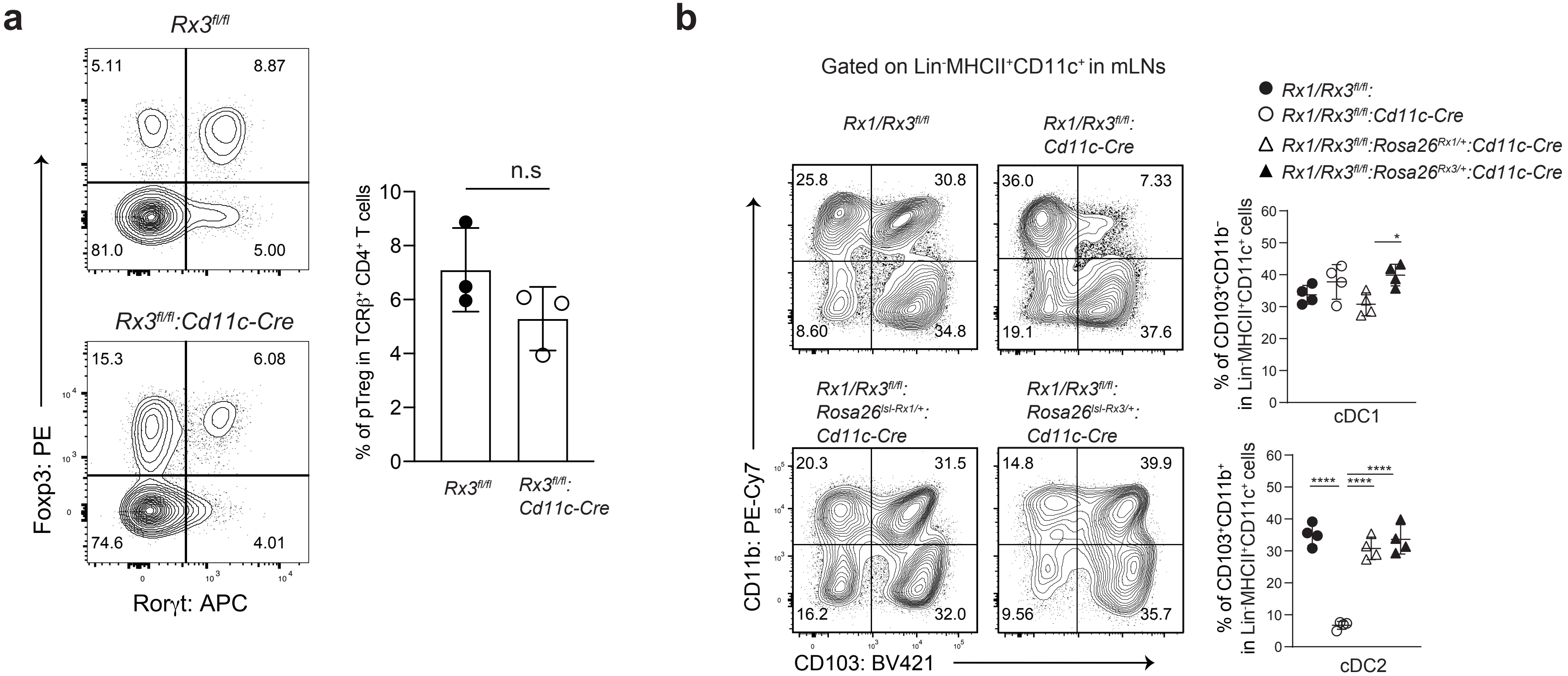


#### Extended Data Figure 3

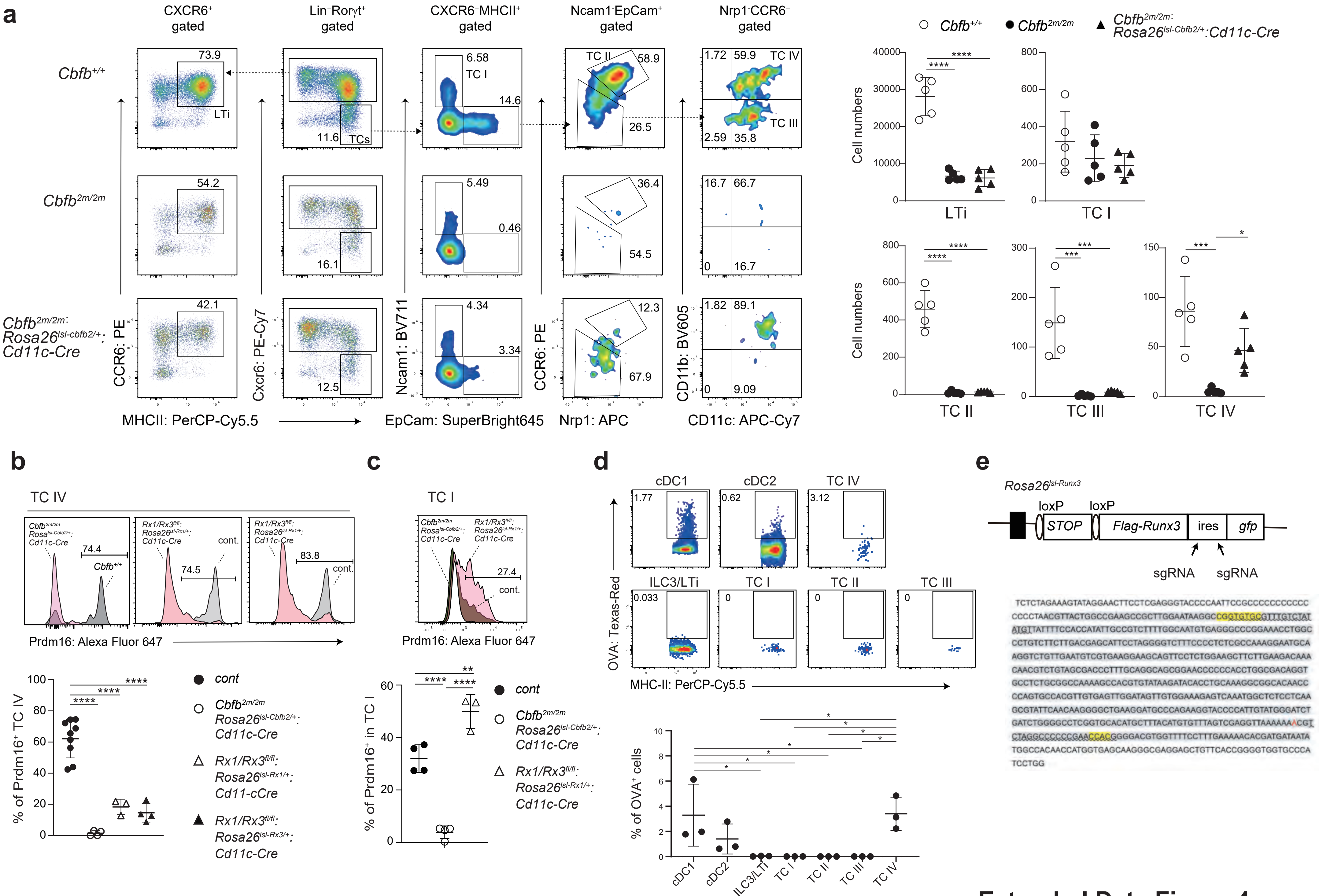


**Extended Data Figure 4**
